# Systematic Evaluation of Somatic Contamination in Germline Genomes

**DOI:** 10.1101/2025.08.21.671542

**Authors:** Xiangwen Ji, Xueke Bai, Guangda He, Kai Yan, Edwin Wang, Yi-Da Tang, Liang Chen, Qinghua Cui

**Affiliations:** Department of Cardiology and Institute of Vascular Medicine, State Key Laboratory of Vascular Homeostasis and Remodeling, Peking University Third Hospital, 49 Huayuanbei Road, Beijing 100191, China; National Clinical Research Center for Cardiovascular Diseases, State Key Laboratory of Cardiovascular Disease, Fuwai Hospital, National Center for Cardiovascular Diseases, Chinese Academy of Medical Sciences and Peking Union Medical College, Beijing, China; Department of Biochemistry and Molecular Biology, Medical Genetics, and Oncology, Cumming School of Medicine, University of Calgary, Calgary, Alberta, Canada; School of Sports Medicine, Wuhan Sports University, No. 461 Luoyu Rd. Hongshan District, Wuhan 430079, Hubei Province, China; Department of Biomedical Informatics, State Key Laboratory of Vascular Homeostasis and Remodeling, School of Basic Medical Sciences, Peking University, 38 Xueyuan Rd, Beijing, 100191, China

## Abstract

Large-scale genomic initiatives like the UK Biobank (UKB) have revolutionized our understanding of human disease. These studies typically assume that blood-derived DNA faithfully reflects an individual’s germline genome. However, this assumption is challenged by somatic mutations arising from processes like clonal hematopoiesis. Although standard bioinformatics pipelines employ variant allele frequency (VAF)-based filtering to mitigate such contamination, the efficacy of these approaches requires systematic evaluation. By systematically analyzing whole-exome sequencing (WES) and whole-genome sequencing (WGS) data from large cohorts, including the UK Biobank, The Cancer Genome Atlas (TCGA), and the 1000 Genomes Project, we revealed critical limitations in current filtering methodologies. We found that the mutational spectrum of rare “germline” variants is highly similar to that of somatic mutations. Furthermore, we uncovered these variants show significant associations with phenotypes such as age, sex, and smoking status, established drivers of somatic mutagenesis. This persistent somatic contamination introduces substantial confounder effects, potentially generating spurious associations and reverse causality in genetic studies. Our work underscores the urgent reconsideration of two fundamental aspects of genomic research: (1) refinement of variant filtering strategies to better distinguish true germline variants from somatic contaminants, and (2) incorporation of somatic mutagenesis factors as essential covariates in study design. Our findings provide a basic framework for improving the accuracy and interpretability of large-scale genomic studies.

## Introduction

The application of whole-exome sequencing (WES) and whole-genome sequencing (WGS) to large-scale population cohorts, such as the UK Biobank (UKB), has significantly advanced our understanding of the genetic basis of complex diseases^1–4^. These massive datasets form the cornerstone of exome-wide association studies (ExWAS), genome-wide association studies (GWAS), and protein quantitative trait loci (pQTL) studies, etc., which aim to identify genetic loci associated with specific traits or diseases by analyzing genetic variation among individuals^5–7^. A central premise of these studies is that blood-derived DNA accurately represents an individual’s germline (i.e., constitutional) genome.

However, this foundational assumption is challenged by the phenomenon of clonal hematopoiesis (CH), a biological process strongly associated with aging^8,9^. In CH, hematopoietic stem cells acquire somatic mutations that confer a selective advantage, leading to their clonal expansion^10^. Consequently, when whole blood is used as the DNA source in WES or WGS studies, these somatic mutations will be admixed into the DNA samples for sequencing. This admixture then introduces a form of “biological noise” that acts as a potential confounder in GWAS, which are designed to identify germline variants. When these somatic mutations, often referred to as CH of indeterminate potential (CHIP), are erroneously identified as germline variants, they will lead to wrong findings in association analyses.

To address this challenge, researchers commonly employ filtering strategies based on variant allele frequency (VAF) to distinguish between germline and somatic variants. The rationale behind this approach is that a true heterozygous germline variant is expected to have a VAF of approximately 50%, whereas a homozygous germline variant will have a VAF of 100%. In contrast, the VAF of a CHIP mutation is typically substantially lower than 50%^11^. Therefore, bioinformatics pipelines^7,12^ typically set a VAF threshold (e.g., ≥20%) and/or utilize a binomial test to remove variants with a VAF significantly below 50%, thereby enriching for high-confidence germline variants. Additionally, metrics such as sequencing depth are considered standard practice to effectively exclude somatic interference and purify the germline signal^7,13^. Although current variant filtering methods are intuitively designed and widely adopted, our study reveals their critical limitation: they fail to fully purge somatic mutational contamination from germline datasets. By systematically analyzing WES and WGS data from large cohorts, including the UK Biobank, The Cancer Genome Atlas (TCGA), and the 1000 Genomes Project, we identified persistent somatic mutational signatures classified as germline, particularly within rare variant subsets. Strikingly, these ostensibly germline variants exhibit molecular profiles characteristic of somatic mutagenesis, suggesting substantial admixture between true germline variants and somatic mosaic events. Such misclassification poses a serious analytical challenge, as residual somatic variants masquerading as germline variants could introduce false-positive associations in genetic studies, ultimately distorting our interpretation of disease etiology. Our findings underscore the imperative to rigorously quantify and account for somatic contamination in germline genomic datasets. Addressing this underappreciated source of bias is essential for enhancing the precision and reproducibility of large-scale genomic studies.

## Methods

### Collection of Germline and Somatic Mutation Data

The UKB stands as a large-scale, prospective cohort study and a comprehensive global biomedical resource. It houses an extensive collection of genetic, proteomic, and phenotypic data from its participants. Between 2006 and 2010, a total of 502,156 individuals, aged 40-69 at baseline, were enrolled from 22 different centers across the UK. For the UKB WES cohort, exomes were captured using the IDT xGen Exome Research Panel v1.0 with supplemental probes and sequenced with 75 bp paired-end reads on the Illumina NovaSeq 6000 platform^1^. The sequencing data were processed through the Original Quality Functional Equivalent (OQFE) pipeline, and germline variants were subsequently called using DeepVariant^14^. Finally, variant information was available for 469,881 participants. This study utilized data from the UKB resource under approved application number 87841.

In addition to the UKB WES cohort, we applied for access to the germline mutation data of 10,389 TCGA patients from a previous study^15^. Germline variant data from the 1000 Genomes Project^16^ Phase 1 (release: 20110521) and Phase 3 (release: 20130502), based on whole-genome sequencing (WGS), were downloaded from the official FTP site (https://ftp.1000genomes.ebi.ac.uk/vol1/ftp/release/).

The Catalogue of Somatic Mutations in Cancer (COSMIC, https://cancer.sanger.ac.uk/cosmic) is the most comprehensive and detailed resource for somatic mutations in human cancer^17^. We downloaded frequency data for coding mutations in haematopoietic and lymphoid tissues from COSMIC v99 (reference genome: GRCh38).

### Germline Variant Filtering

To filter germline variants, we adopted a strategy similar to that used in our previous work^13^. First, a minimum sequencing depth of 20 was required to ensure sequencing reliability. The VAF was calculated as the ratio of variant allele depth (AD) to total depth (DP), i.e., VAF=AD/DP. To exclude potential somatic mutations, such as those associated with CHIP, which typically present with low VAF, we fitted the VAF distribution of all variants using a Gaussian Mixture Model (GMM). In previous study, we defined variants with a VAF between 0.422 and 0.54 or ≥ 0.9 as high-confidence germline mutations.

Furthermore, to assess the robustness of our results, we also tested five alternative filtering strategies: 1) DP ≥ 10, and either 0.422 ≤ VAF ≤ 0.54 or VAF ≥ 0.9; 2) DP ≥ 20, VAF ≥ 0.2, and binomial test p-value ≥ 1×10^-6^ for heterozygous; 3) DP ≥ 10, VAF ≥ 0.2, and binomial test p-value ≥ 1×10^-6^ for heterozygous; 4) DP ≥ 20 and AD ≥ 5; 5) DP ≥ 10 and AD ≥ 3. Among these, using a binomial test to retain variants whose allele frequencies are not strongly significantly (p ≥ 1×10^-6^) different from a true heterozygous state (VAF=50%) is a common strategy in UKB-based studies to exclude somatic mosaicism^7,12^. Based on the filtering results, our primary strategy proved to be the most stringent, yielding the lowest number of distinct variants (1,447,265) and total variant calls (9,571,462,124).

As AD and DP information was not available for the 1000 Genomes Project data, we could not apply the aforementioned filters. However, these data have undergone rigorous quality control by the 1000 Genomes Project Consortium^16^, lending confidence to the reliability of the variant calls.

### Germline Genome Pattern (GGP) Analysis

Our GGP analysis followed a procedure similar to SigProfiler^18^ and our previous study^13^. Specifically, for each single base substitution (SBS), we identified the base substitution and its immediate 5’ and 3’ flanking nucleotides. This defined a trinucleotide context, resulting in 96 possible mutation types for pyrimidine (C/T) substitutions (4 types of 5’ base × 6 types of substitution × 4 types of 3’ base). Substitutions involving purines (G/A) were reverse complemented to be represented by their pyrimidine-based equivalents.

For each sample, all SBS germline variants were classified into one of the 96 trinucleotide contexts, and the counts for each context were tallied. For a cohort of *N* samples, this process generates a 96 × *N* matrix, denoted as *X*. Using a Non-negative Matrix Factorization (NMF) algorithm^19^, matrix *X* can be decomposed into two matrices: a 96 × *k* signature matrix *W* and a *k* × *N* activity matrix *H*, such that:

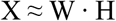

The matrices can be scaled such that each column of *W* sums to 1, allowing it to be interpreted as a probability distribution representing a distinct mutational signature. Consequently, each column in matrix *H* contains *k* values, representing the activities of the *k* signatures (e.g., GGP A, GGP B, etc.) in the corresponding sample. For new data, the signature activities can be determined by fitting a new activity matrix *H* while keeping the signature matrix *W* fixed.

To determine the optimal number of signatures, *k*, we iterated from 1 to 25. For each *k*, we performed a resampling procedure for 100 times. In each resampling iteration, a new mutation count matrix was generated by sampling from multinomial distributions based on each sample’s original mutation profile. NMF was applied to each resampled matrix to extract 100 sets of *k* signatures. These 100×k signatures were then clustered into *k* groups using K-means clustering. The Hungarian algorithm^20^ was used to match signatures across iterations, minimizing cosine distances to cluster centroids. The silhouette coefficient (using cosine distance) was used to evaluate clustering quality. For *k*=1, the coefficient was set as 1. We selected the largest *k* for which the coefficient remained above 0.7. The final *k* de novo signatures were defined by averaging the 100 signatures within each cluster. These computations were performed using Python (v3.13). NMF was implemented with torchnmf (v0.3.5), K-means clustering and silhouette coefficient were implemented with scikit-learn^21^ (v1.7.0), the Hungarian algorithm was implemented with SciPy^22^ (v1.15.3), and multinomial sampling was implemented with NumPy^23^ (v2.3.0).

After obtaining the de novo signatures, we decomposed them into known COSMIC mutational signatures (v3.4) using SigProfilerAssignment^24^ (v0.2.3). A non-negative least squares (NNLS) approach, with a regularization penalty to prevent overfitting, was used to represent each de novo signature as a linear combination of known signatures (requiring cosine similarity > 0.8). We excluded signatures associated with DNA repair and chemotherapy from candidate signatures as they were not relevant to this study. The algorithm then refits the activity matrix *H* using the established COSMIC signature matrix *W*. The values in the refitted *H* matrix are constrained to integers, representing the estimated number of mutations attributed to each signature. The mutation burden defined as number of mutations per megabase (mut/Mb) was calculated by assuming that an average of 39 Mb of the exome has sufficient coverage, based on UKB sequencing protocols^1^.

### GWAS Data Collection

We downloaded published GWAS summary statistics (v1.0.e114) from the GWAS Catalog^25^ (https://www.ebi.ac.uk/gwas/). The reference and variant alleles of each association were annotated using dbSNP^26^ (v151, GRCh38), which was downloaded from the NCBI FTP server (https://ftp.ncbi.nlm.nih.gov/snp/organisms/). These variants were subsequently categorized into six substitution types (C>A, C>G, C>T, T>A, T>C, and T>G) and other categories (e.g., insertions and deletions). To reduce the error in calculating the percentage of C>A mutations, only phenotypes with at least 50 associated loci were retained for further analysis. Consequently, a total of 382,636 loci associated with 1,668 phenotypes were included in the final analysis.

Next, we manually selected phenotypes that exhibited a strong positive correlation with smoking based on the following criteria: (1) include phenotypes that directly define smoking status (e.g., “ever vs. never smokers”) or quantify smoking intensity and frequency (e.g., “cigarettes smoked per day”); (2) include respiratory diseases strongly associated with smoking, including lung adenocarcinoma, chronic obstructive pulmonary disease (COPD), pulmonary fibrosis, etc.; and (3) exclude phenotypes that had already been adjusted for smoking status. By doing so, 24 phenotypes met these criteria (Supplementary Table S5).

### Statistical Analysis

Correlations between continuous variables were assessed using the two-sided Spearman’s rank correlation. The difference between two groups of continuous variables was evaluated using the two-sided Wilcoxon rank-sum test. Furthermore, multiple linear regression models were used to evaluate the contribution of phenotypes to either the total mutation burden or the activity of specific mutational signatures, providing coefficients, 95% confidence intervals (CIs), and p-values. The following UKB phenotype fields were used as covariates: age (field 21003), sex (field 31), and smoking status (field 20116). All statistical analyses were performed using the SciPy^22^ (v1.15.3) and statsmodels^27^ (v0.14.4) Python packages.

## Results

### The Frequency-Dependent Spectral Characteristics of Germline and Somatic Mutations

Somatic mutations, due to their relatively random distribution across the genome, are expected to recur at the same genomic position with a low frequency. Consequently, low-frequency germline mutations are more likely to be admixed with de novo somatic mutations. Therefore, we compared the relationship between the spectral characteristics and the number of occurrences for germline mutations derived from blood WES (from UKB) and somatic mutations in haematopoietic and lymphoid tissues (from COSMIC). The result demonstrates that these two classes of mutations demonstrated markedly different frequency dependencies (Figure 1).

**Figure 1.**
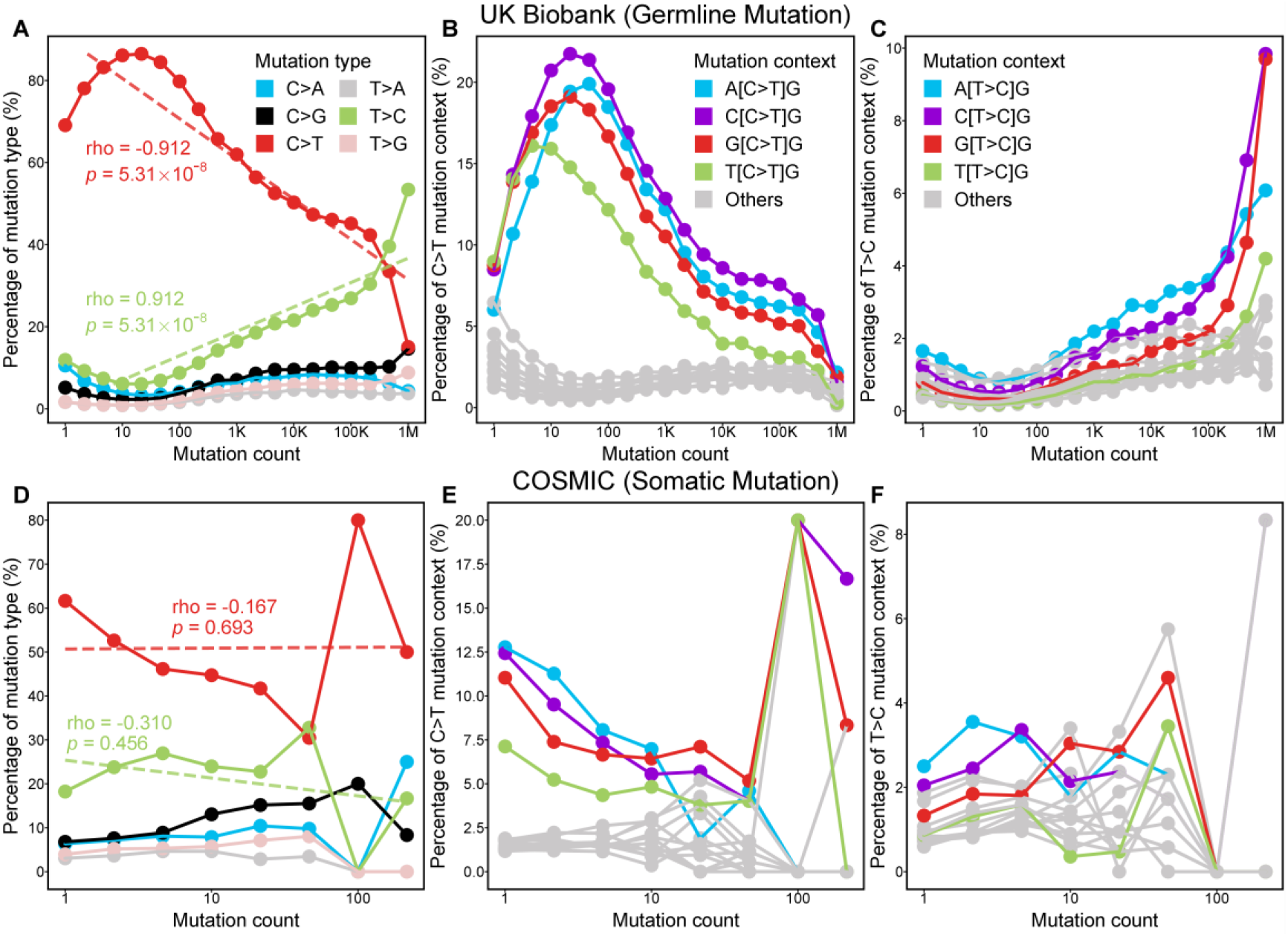
Frequency dependence of mutational spectra in germline vs. somatic mutations. (A-C) Analysis of germline mutations from UK Biobank (UKB). (D-F) Analysis of somatic mutations from the COSMIC database, sharing the same color legends with (A-C), respectively. (A, D) The relationship between the percentage of the six base substitution types (C>A, C>G, C>T, T>A, T>C, T>G) and the mutation count. The Spearman’s rank correlation coefficient (rho) and p-value for the correlation between the percentage of C>T (red) and T>C (green) mutations and the mutation count are shown. (B, E) The percentage of C>T mutations within various trinucleotide contexts, plotted against the mutation count. (C, F) The percentage of T>C mutations within various trinucleotide contexts, plotted against the mutation count. The x-axis is represented on a logarithmic scale.

For germline mutations, we observed a strong correlation between the proportion of certain mutation types and their counts. Specifically, the relative abundance of C>T (G>A) transitions showed a significant negative correlation with mutation count (Figure 1A, Spearman’s ρ=-0.912, *p*=5.31×10^-8^). For mutations with counts below 100, C>T mutations constituted nearly 80% of the total. A closer examination of the trinucleotide context of C>T mutations revealed a higher proportion of the four contexts with a downstream guanine (NpCpG) (Figure 1B). Mutations arising from the deamination of 5-methylcytosine (5-mC) at CpG dinucleotides, which have a high spontaneous mutation rate, are recognized as an age-related “clock-like” signature in somatic mutation studies^28,29^. Conversely, T>C (A>G) transitions represented the second most dominant mutation type, and their relative abundance was significantly positively correlated with mutation count (Figure 1A, Spearman’s ρ=0.912, *p*=5.31×10^-8^), eventually surpassing C>T mutations in abundance at the high-frequency end of the curve. Similarly, the four mutational contexts with a downstream guanine were the primary constituents of T>C mutations (Figure 1C). This may be an artifact resulting from differences between the human reference genome and the common genotype in the UK population, as a T>C mutation for most individuals can be interpreted as a C>T mutation for a minority. We further investigated blood WES data from TCGA and WGS of blood-derived lymphoblastoid cell lines (LCLs) from Phases 1 and 3 of the 1000 Genomes Project. As a result, similar features were observed (Supplementary Figure S1), suggesting this pattern is independent of population, sequencing technology, and reference genome version. Moreover, this phenomenon remained robust under different germline variant filtering criteria (Supplementary Figure S2). It should be noted that we have employed a stringent filtering strategy that yields a minimal number of variants (Supplementary Figure S2A). Using another common filtering method like the binomial test would result in a larger set of retained variants (Supplementary Figure S2C). Interestingly, for rare variants, there was an abnormal decrease in the proportion of C>T mutations, accompanied by an increase in the proportion of mutation types dominated by C>A (G>T) transversions. This increase was more pronounced under lenient filtering criteria, where C>A transversions could even surpass T>C mutations, the original second-most abundant type (Supplementary Figure S2F). This suggests that rare variants may be contaminated with somatic mutations arising from other etiological factors. For instance, polycyclic aromatic hydrocarbons in tobacco smoke can form adducts with bases in CpG islands, a known cause of C>A transversions in smokers’ genomes^30,31^.

In contrast, we did not observe such clear patterns among somatic mutations. Although C>T and T>C mutations still represented the two most dominant mutation types, their relative abundances showed no statistically significant correlation with their counts in the COSMIC database (Figure 1D; C>T: ρ=-0.167, *p*=0.693; T>C: ρ=-0.310, *p*=0.456). The entire mutational spectrum appeared more random and noisier across different frequency bins. Likewise, at the trinucleotide context level, the distribution of somatic mutations did not exhibit any uniform regularity with respect to frequency (Figures 1E, 1F). As statistical power is affected by the number of data points, we repeated the experiment with different logarithmic scale bin sizes. The results confirmed that the above statistical outcomes are not influenced by bin size. The significant changes in the UKB germline mutation spectrum remained observable with larger bin sizes and fewer data points (Supplementary Figure S3A), while the non-significant changes in the COSMIC somatic mutation spectrum remained non-significant even with smaller bin sizes and more data points (Supplementary Figure S3B). These findings indicate that somatic mutations have a stable frequency distribution, which more closely resembles that of low-frequency germline mutations. This offers a new perspective on how mutational processes at the levels of natural selection and somatic development shape the evolution of the human genome.

### Mutational Signatures Reveal Contextual Features of Germline Mutations

To investigate the somatic-like features embedded within the germline mutation spectrum, we analyzed the SBS signatures in all variants and variants with allele frequency (AF) ≤ 0.1% using GGP analysis (Figure 2). In our previous studies^13,32^, GGP successfully identified clinically relevant signatures in cancer and COVID-19 patients. For rare variants, three de novo GGPs were identified (Figure 2A-C, Supplementary Table S1, S2). However, all could be explained by different proportions of classic somatic mutational signatures, SBS1 and SBS5. SBS1^28^ displays a high frequency of C>T transitions within NpCpG contexts (Figure 2E) and SBS5^33^ contains a broad components involving nearly all types of SBSs (Figure 2F). Although SBS5 was identified early, the mutational process it represents remains unclear, with only statistical associations to aging, smoking, and deficiencies in nucleotide excision repair^34^. The detection of a similar signature in the germline genome suggests that SBS5 may represent an average mutational pattern arising from various mutational processes with weak base specificity.

**Figure 2.**
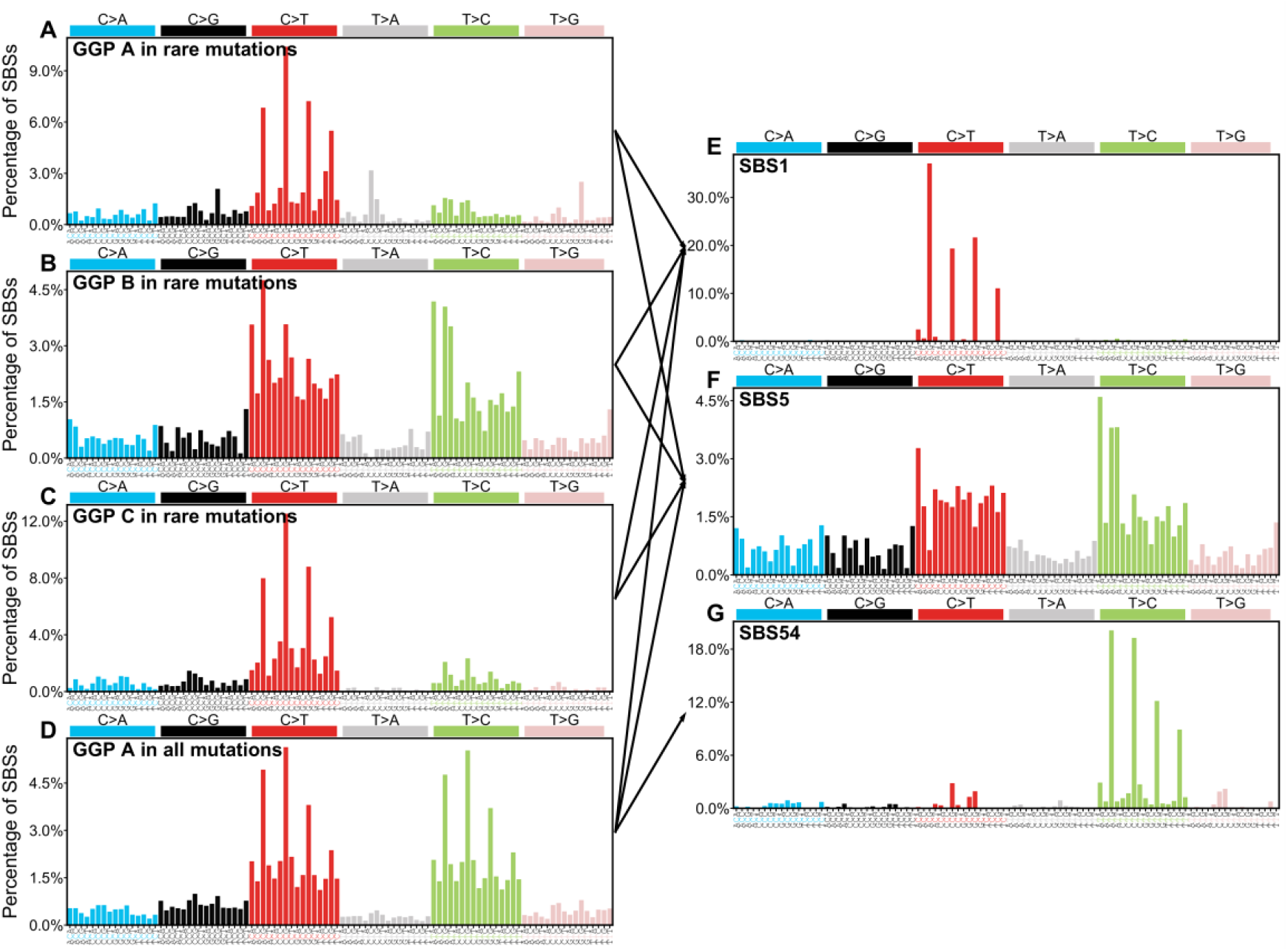
Germline Genome Patterns (GGPs) and their matching to known COSMIC signatures. (A-C) Three de novo GGPs extracted from rare mutations. (D) One de novo GGP extracted from all mutations. (E-G) Reference COSMIC mutational signatures corresponding to the patterns in (A-D). The x-axis for all panels displays the 96 trinucleotide contexts of single base substitutions, which are defined by six substitution types (C>A, C>G, C>T, T>A, T>C, T>G, shown in distinct colors) and their 5’ and 3’ flanking nucleotides. The y-axis represents the percentage of each context. Arrows connecting the left and right indicate the relationship between the de novo GGPs and reference signatures, as determined by non-negative least squares optimization.

For the entire set of mutations, only a single GGP was extracted (Figure 2D, Supplementary Table S3, S4). This is likely because the general background genomic pattern of the population is dominant, masking the effects of other mutational etiologies. This single signature could be decomposed into three COSMIC SBS signatures: SBS1, SBS5, and SBS54. SBS54 is characterized by T>C mutations in NTG contexts (Figure 2G) and is often considered an artifact caused by germline mutations in somatic signature studies^34^. As discussed above, this is likely due to an inconsistency between the reference genome and the population’s genotype. Given the lower proportion of T>C mutations among rare variants, the absence of SBS54 in rare variants is then reasonable.

### Germline Mutation Burden is Associated with Mutagenic Etiologies

To evaluate the contamination of somatic mutations in germline mutation calls, we used a multiple linear regression model to explore the relationships between germline mutation burden and age, sex, and smoking status (previous and current). To eliminate confounding effects of ethnicity, only individuals of White ethnicity (self-reported ethnicity, UKB field 21000), who constitute 94.1% of the UKB WES cohort, were included. Furthermore, we restricted the analysis to autosomal mutation burden to avoid sex-related biases. The basic statistics for the variables are presented in Table 1.

**Table 1.** Characteristics and mutation burden of the study cohort from UK Biobank.

|  | Category | Unit | Statistics |
| --- | --- | --- | --- |
| **n** | - | - | 441,213 |
| **Age, mean (std)** | - | year | 56.5 (8.09) |
| **Sex, n (%)** | Female | - | 239,412 (54.3) |
|  | Male | - | 201,801 (45.7) |
| **Smoking status, n (%)** | Never | - | 237,998 (53.9) |
|  | Previous | - | 157,024 (35.6) |
|  | Current | - | 46,191 (10.5) |
| **Total burden, mean (std)** | Total mutation | mut/Mb | 809 (104) |
|  | Rare mutation | mut/Mb | 7.79 (2.12) |
| **SBS1 burden, mean (std)** | Total mutation | mut/Mb | 120 (13.0) |
|  | Rare mutation | mut/Mb | 1.93 (0.491) |
| **SBS5 burden, mean (std)** | Total mutation | mut/Mb | 527 (79.0) |
|  | Rare mutation | mut/Mb | 5.86 (1.73) |
| **SBS54 burden, mean (std)** | Total mutation | mut/Mb | 162 (13.8) |

Abbreviations: std, standard deviation with delta degree(s) of freedom=1; mut/Mb, mutations per megabase; SBS, single base substitution. SBS1, SBS5, and SBS54 refer to specific mutational signatures defined by COSMIC database.

The results showed that for the total germline mutation burden (Figure 3A), current smoking status was estimated to contribute an additional 1.29 mut/Mb (95% CI: 0.253-2.33, *p*=0.0148).

**Figure 3.**
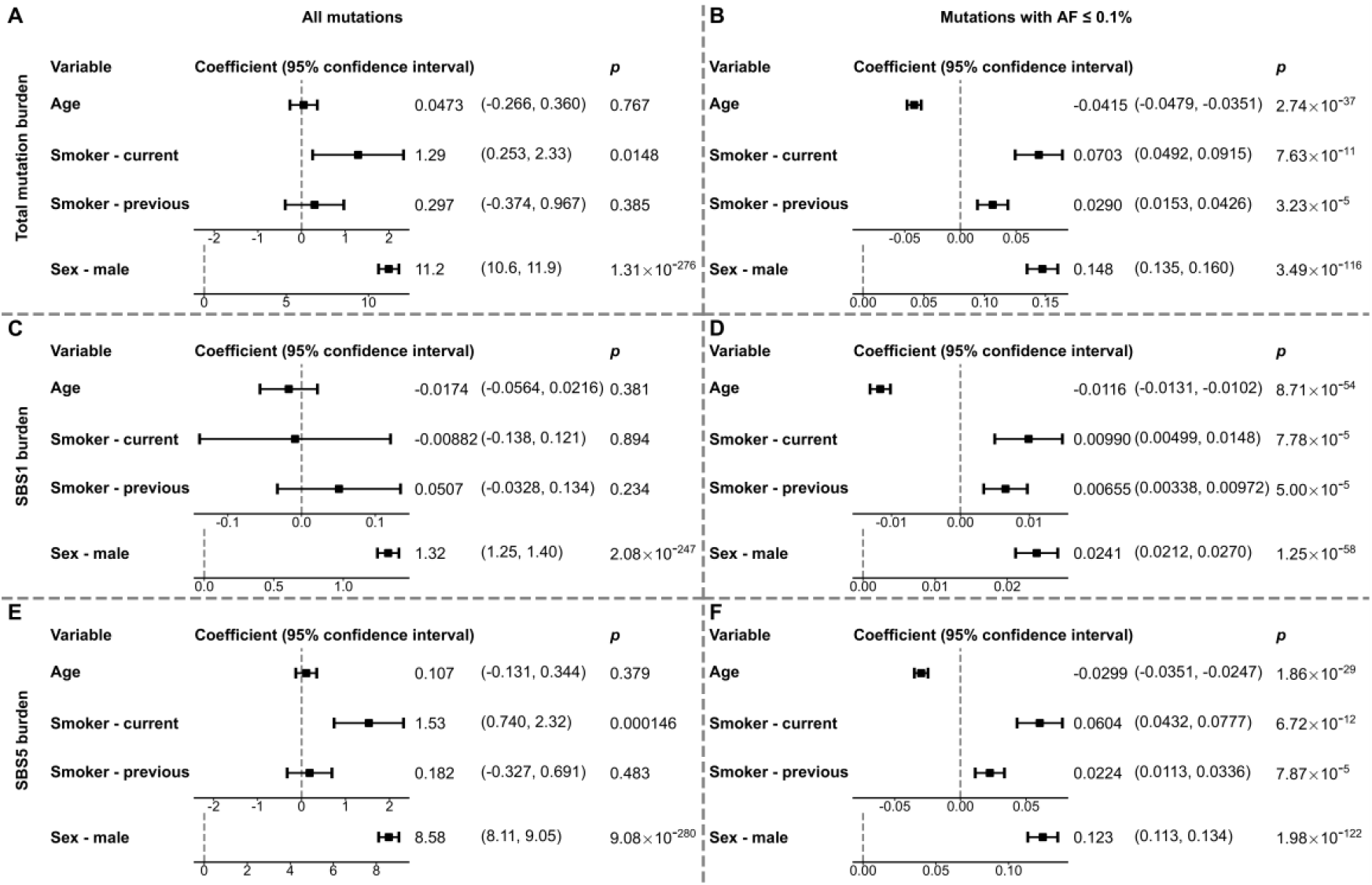
Multivariable analysis of mutagenic factors influencing germline mutation burdens. This figure displays forest plots from multivariable linear regression models, examining the associations of age, smoking status, and sex with (A, B) total germline mutation burden, (C, D) SBS1-related germline mutation burden, and (E, F) SBS5-related germline mutation burden. The left panels (A, C, E) present the results from analyses using all mutations. The right panels (B, D, F) show the results from analyses restricted to mutations with allele frequencies ≤ 0.1%. The square points represent the estimates of the regression coefficients, and the horizontal lines indicate the 95% confidence intervals (CIs). For smoking status, “never smoking” was used as the reference group, while “female” is the reference for the sex variable. The coefficients, 95% CIs, and p-values for each variable are listed to the right of each panel. n=441,213.

More strikingly, males were found to accumulate 11.2 more mut/Mb than females (95% CI: 10.6-11.9, *p*=1.31×10^-276^), possibly reflecting different rates of mutation accumulation due to differences in DNA repair and hormone levels between sexes^35,36^. As the median tumor mutation burden for many types of cancer is less than 10 mut/Mb^37,38^, these findings could present a significant impact.

Notably, the burden of rare variants exhibited even more significant associations with these phenotypes (Figure 3B). In addition to the burden increase of 0.0703 mut/Mb for current smokers (95% CI: 0.0492-0.0915, *p*=7.63×10^-11^), a significant increase was also observed for previous smokers (0.0290 mut/Mb; 95% CI: 0.0153-0.0426, *p*=3.23×10^-5^). This indicates that the damage of smoking to germline genome are long-lasting and persist even after cessation. Considering the proportion of effects, the impact of current smoking accounted for 0.159% of the total mutation burden but 0.903% of the rare mutation burden, suggesting a higher proportion of somatic contamination among rare variants.

A paradoxical observation was a significant negative correlation between age and mutation burden (Figure 3B), where each standard deviation (std, 8.09 years) increase in age was associated with a decrease of 0.0415 mut/Mb (95% CI: -0.0479 to -0.0351, *p*=2.74×10^-37^). This contradicts the generally accepted positive correlation between age and mutational burden in somatic studies^8,34^. We hypothesize that our stringent variant filtering strategy may have filtered out more potential somatic mutations and rare mutations in older individuals, causing the remaining variants to exhibit a negative correlation with age. Therefore, the relationship between the burden of mutation with ultra-small AF (≤ 0.01%) and age was further investigated, and a more significant negative correlation was observed (-0.0163 mut/Mb with 95% CI: - 0.0187 to -0.0139, *p*=3.33×10^-40^). This would also imply that our estimates of the effects of other mutagenic factors are conservative, and their actual impact on mutation burden may be more severe.

The burdens of mutations attributed to SBS1 and SBS5 showed results similar to the total burden for both all and rare variants (Figure 3C-F). As smoking primarily induces C>A mutations with weaker effects on other mutation types, the association between smoking and the C>T-dominant SBS1 burden was weaker than its association with SBS5, although it remained significant among rare mutations. In summary, our stratified analysis reveals the influence of various factors on germline mutation burden, challenging the conventional view of causality from the germline genome to phenotype.

### Spurious Germline Mutations May Lead to False-Positive GWAS Results

Smoking, a major risk factor for multiple cancers and respiratory diseases, is widely recognized for inducing a higher proportion of C>A transversions in somatic genomes^31,39^. Given our finding that smoking also generates a considerable mutational burden in the germline genome, we wondered whether GWAS of smoking-related phenotypes have been affected.

Therefore, we collected 382,636 association loci for 1,668 phenotypes and calculated the proportion of C>A mutations at the associated loci for each phenotype (Supplementary Table S5). Among these, we focused on 24 phenotypes due to their strong positive correlation with smoking. Notably, 18 (75%) of these smoking-related phenotypes, such as smoking quantity (cigarettes per day, based on multi-trait analysis of GWAS [MTAG]), idiopathic pulmonary fibrosis, and smoking status (ever vs. never smoker), ranked in the top 50% for their percentage of associated C>A mutations (Figure 4A). Statistical analysis revealed that the percentage of C>A transversions among mutations associated with smoking-related phenotypes was significantly higher than in those associated with other phenotypes (Figure 4B, *p*=0.0469, two-sided Wilcoxon rank-sum test). Our findings suggest that artifacts from smoking-induced germline mutations may have already biased previously published GWAS results. These mutations and the smoking-associated phenotypes are likely common consequences of smoking, creating a fallacious causal link from germline polymorphism to phenotype that could misdirect subsequent research based on these findings.

**Figure 4.**
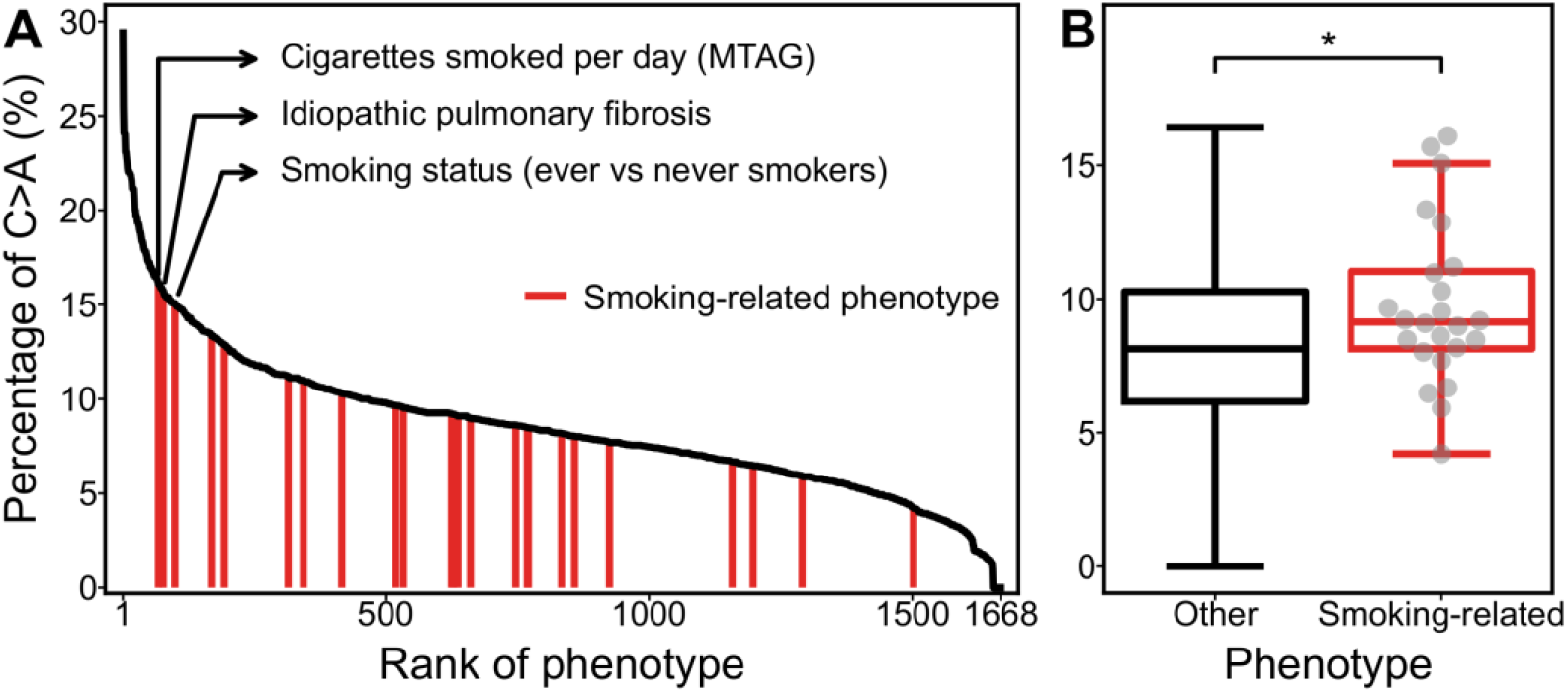
Loci associated with smoking-related phenotypes exhibit a higher proportion of C>A mutations. (A) Associated risk loci for 1,668 phenotypes are sorted in descending order by the percentage of C>A mutations. Phenotypes manually annotated as smoking-related are marked by red vertical lines. Arrows highlight the 3 smoking-related phenotypes with the highest C>A mutation percentages. (B) Box plot comparing the percentage of C>A mutations between the smoking-related phenotype group (red box, n = 24) and all other phenotypes (black box, n = 1,644). The center line of each box represents the median; the lower and upper hinges of the box correspond to the first and third quartiles, respectively. The whiskers extend to the furthest data points within 1.5 times the interquartile range (IQR). The dots in the smoking-related group represent the values for each individual phenotype. \**p*<0.05 by two-sided Wilcoxon rank-sum test.

## Discussion

De novo germline mutations in a new generation originate from somatic mutations within the parental germ cells. In this study, we observed that the mutational profile of low-frequency germline variants is highly consistent with the characteristics of somatic mutations found in cancer genomics. We successfully decomposed the classic somatic mutational signatures, SBS1 and SBS5, from the spectrum of germline mutations. As “clock-like” signatures, SBS1 and SBS5 are detectable in nearly all tumor types and even in normal tissues^34,40^. The striking similarity between germline and somatic mutational signatures suggests a unified principle underlying species evolution at the population level and somatic clonal evolution at the individual level.

Previous research has established that CHIP is a common age-related phenomenon, and its associated mutations (e.g., in genes such as DNMT3A and TET2) are risk factors for cardiovascular disease and hematologic malignancies^41–43^. However, the influence of CHIP signals has been rarely considered in GWAS. Our analysis reveals theoretically implausible yet significant associations between germline mutation burden and age, sex, and smoking status, particularly for rare mutations. These findings challenge the current paradigm of genetic association studies and Mendelian randomization studies by highlighting the potential for CHIP-mediated false-positive results and inverting cause and effect. For instance, smoking induces somatic mutations in hematopoietic stem cells and is also a cause of smoking-related diseases such as COPD and lung cancer. This can lead to spurious associations in GWAS between smoking-induced pseudo “germline” variants and the disease. In our prior work^13^, we identified seven cancer GGPs (CGGPs) from germline genomes in TCGA. One of these, CGGP E, exhibited a robust and significant positive correlation with the somatic mutational signature SBS4, a well-established signature associated with tobacco smoking^30^, independent of cancer type, sex, and age. Assuming that the germline genome will not change afterbirth, we initially interpreted this as evidence of the germline genome’s influence on an individual’s susceptibility to smoking-related diseases. From our current perspective, however, these findings suggest that smoking may have imposed a wide mutation burden on the “germline” genome, with both CGGP E and SBS4 being joint consequences of tobacco exposure.

A seemingly contradictory finding in this study is the negative correlation between age and the mutation burden after filtering. As age increases, the number of CHIP mutations rises substantially. Consequently, any filtering strategy designed to remove CHIP will inevitably exclude a greater number of variants from older individuals compared to younger ones. This filtering bias creates an artifactual negative association. Furthermore, this observation implies that the estimated impact of other factors, such as smoking, may also be severely underestimated.

This research reveals the stubborn contamination of somatic mutations within germline genomic data from large-scale UKB WES cohort. We should recognize the limitations of current filtering strategies and reconsider the definition and operational standards of “germline” in studies utilizing blood-derived DNA. For example, machine learning models could be developed to more accurately estimate the probability of a variant being of germline or somatic origin by integrating multi-dimensional features, including VAF, trinucleotide context, sequencing depth, strand bias, and known CHIP drivers. Moreover, incorporating CHIP status and exposure to mutagens like tobacco smoke as covariates in statistical models may be an effective strategy to mitigate their confounding effects in association analyses. Nevertheless, this study has its limitations. Our analysis was predominantly based on the White population from the UK, and validation in large-scale cohorts of other ancestries is lacking. Additionally, a long-term, longitudinal study with repeated sequencing is warranted to more precisely assess the dynamics of the germline genome over an individual’s lifespan.

## Data availability

The data used in this study are available from the UKB under specific restrictions. As the data were accessed under license, they are not publicly available. Access to UK Biobank data can be requested through the standard application process (https://www.ukbiobank.ac.uk/registerapply/). The TCGA germline mutation data are available according to the authorization procedure described on this website: http://isb-cancer-genomics-cloud.readthedocs.io/en/latest/sections/webapp/Gaining-Access-To-Contolled-Access-Data.html. The 1000 Genomes Project data are available from https://ftp.1000genomes.ebi.ac.uk/vol1/ftp/release/. The frequencies of somatic mutations are available from COSMIC, https://cancer.sanger.ac.uk/cosmic. The GWAS associations are available from GWAS Catalog, https://www.ebi.ac.uk/gwas/, and the variants can be annotated using dbSNP, https://ftp.ncbi.nlm.nih.gov/snp/organisms/. All data supporting the findings of this study are included in the article and supplementary materials and can be obtained from the corresponding author upon request.

## Acknowledgements

This study was supported by the grants from the National Natural Science Foundation of China [62025102, 82470373]

## Author contributions

X.J. and Q.C. designed the study. X.J. performed the analysis. X.B. and G.H. took part in the discussion. K.Y. and E.W. took part in running some of the algorithms. X.J. wrote the manuscript and Q.C. edited the manuscript. Y.D.T, L.C. and Q.C. supervised the study.

## References

1. Van Hout, C. V. et al. Exome sequencing and characterization of 49,960 individuals in the UK Biobank. Nature 586, 749–756 (2020).

2. Backman, J. D. et al. Exome sequencing and analysis of 454,787 UK Biobank participants. Nature 599, 628–634 (2021).

3. Halldorsson, B. V. et al. The sequences of 150,119 genomes in the UK Biobank. Nature 607, 732–740 (2022).

4. Burren, O. S. et al. Genetic architecture of telomere length in 462,666 UK Biobank whole-genome sequences. Nat Genet 56, 1832–1840 (2024).

5. Shen, S. et al. A Large-Scale Exome-Wide Association Study Identifies Novel Germline Mutations in Lung Cancer. Am J Respir Crit Care Med 208, 280–289 (2023).

6. Szustakowski, J. D. et al. Advancing human genetics research and drug discovery through exome sequencing of the UK Biobank. Nat Genet 53, 942–948 (2021).

7. Dhindsa, R. S. et al. Rare variant associations with plasma protein levels in the UK Biobank. Nature 622, 339–347 (2023).

8. Wen, S. et al. Comparative analysis of the Mexico City Prospective Study and the UK Biobank identifies ancestry-specific effects on clonal hematopoiesis. Nat Genet 57, 572–582 (2025).

9. Xie, M. et al. Age-related mutations associated with clonal hematopoietic expansion and malignancies. Nat Med 20, 1472–1478 (2014).

10. Natarajan, P., Jaiswal, S. & Kathiresan, S. Clonal Hematopoiesis: Somatic Mutations in Blood Cells and Atherosclerosis. Circ Genom Precis Med 11, e001926 (2018).

11. Genovese, G. et al. Clonal hematopoiesis and blood-cancer risk inferred from blood DNA sequence. N Engl J Med 371, 2477–2487 (2014).

12. Wang, Q. et al. Rare variant contribution to human disease in 281,104 UK Biobank exomes. Nature 597, 527–532 (2021).

13. Xu, X. et al. Germline genomic patterns are associated with cancer risk, oncogenic pathways, and clinical outcomes. Sci Adv 6, eaba4905 (2020).

14. Regier, A. A. et al. Functional equivalence of genome sequencing analysis pipelines enables harmonized variant calling across human genetics projects. Nat Commun 9, 4038 (2018).

15. Huang, K.-L. et al. Pathogenic Germline Variants in 10,389 Adult Cancers. Cell 173, 355-370.e14 (2018).

16. 1000 Genomes Project Consortium et al. A global reference for human genetic variation. Nature 526, 68–74 (2015).

17. Tate, J. G. et al. COSMIC: the Catalogue Of Somatic Mutations In Cancer. Nucleic Acids Res 47, D941–D947 (2019).

18. Islam, S. M. A. et al. Uncovering novel mutational signatures by de novo extraction with SigProfilerExtractor. Cell Genom 2, None (2022).

19. Lee, D. D. & Seung, H. S. Learning the parts of objects by non-negative matrix factorization. Nature 401, 788–791 (1999).

20. Kuhn, H. W. The Hungarian method for the assignment problem. Naval Research Logistics Quarterly 2, 83–97 (1955).

21. Pedregosa, F. et al. Scikit-learn: Machine Learning in Python. J Mach Learn Res 12, 2825–2830 (2011).

22. Virtanen, P. et al. SciPy 1.0: Fundamental Algorithms for Scientific Computing in Python. Nature Methods 17, 261–272 (2020).

23. Harris, C. R. et al. Array programming with NumPy. Nature 585, 357–362 (2020).

24. Díaz-Gay, M. et al. Assigning mutational signatures to individual samples and individual somatic mutations with SigProfilerAssignment. Bioinformatics 39, btad756 (2023).

25. Sollis, E. et al. The NHGRI-EBI GWAS Catalog: knowledgebase and deposition resource. Nucleic Acids Res 51, D977–D985 (2023).

26. Sherry, S. T. et al. dbSNP: the NCBI database of genetic variation. Nucleic Acids Res 29, 308–311 (2001).

27. Seabold, S. & Perktold, J. statsmodels: Econometric and statistical modeling with python. in 9th Python in Science Conference.

28. Alexandrov, L. B. et al. Clock-like mutational processes in human somatic cells. Nat Genet 47, 1402–1407 (2015).

29. Pfeifer, G. P. Mutagenesis at methylated CpG sequences. Curr Top Microbiol Immunol 301, 259–281 (2006).

30. Nik-Zainal, S. et al. The genome as a record of environmental exposure. Mutagenesis 30, 763–770 (2015).

31. Pfeifer, G. P. et al. Tobacco smoke carcinogens, DNA damage and p53 mutations in smoking-associated cancers. Oncogene 21, 7435–7451 (2002).

32. Zhang, N. et al. Genomic Patterns are Associated with Different Sequelae of Patients with Long-Term COVID-19. Adv Sci (Weinh) 12, e2407342 (2025).

33. Alexandrov, L. B. et al. Signatures of mutational processes in human cancer. Nature 500, 415–421 (2013).

34. Alexandrov, L. B. et al. The repertoire of mutational signatures in human cancer. Nature 578, 94–101 (2020).

35. Bergeron, L. A. et al. Evolution of the germline mutation rate across vertebrates. Nature 615, 285–291 (2023).

36. Wengner, A. M., Scholz, A. & Haendler, B. Targeting DNA Damage Response in Prostate and Breast Cancer. Int J Mol Sci 21, 8273 (2020).

37. Wu, H.-X. et al. Tumor mutational and indel burden: a systematic pan-cancer evaluation as prognostic biomarkers. Ann Transl Med 7, 640 (2019).

38. Jeong, A.-R. et al. Higher tumor mutational burden and PD-L1 expression correlate with shorter survival in hematologic malignancies. Ther Adv Med Oncol 16, 17588359241273053 (2024).

39. Olivier, M., Hollstein, M. & Hainaut, P. TP53 mutations in human cancers: origins, consequences, and clinical use. Cold Spring Harb Perspect Biol 2, a001008 (2010).

40. Moore, L. et al. The mutational landscape of human somatic and germline cells. Nature 597, 381–386 (2021).

41. Bernstein, N. et al. Analysis of somatic mutations in whole blood from 200,618 individuals identifies pervasive positive selection and novel drivers of clonal hematopoiesis. Nat Genet 56, 1147–1155 (2024).

42. Jaiswal, S. et al. Age-related clonal hematopoiesis associated with adverse outcomes. N Engl J Med 371, 2488–2498 (2014).

43. Genovese, G. et al. Clonal hematopoiesis and blood-cancer risk inferred from blood DNA sequence. N Engl J Med 371, 2477–2487 (2014).

