## Supplementary Figure for "Systematic Evaluation of Somatic Contamination in Germline Genomes"

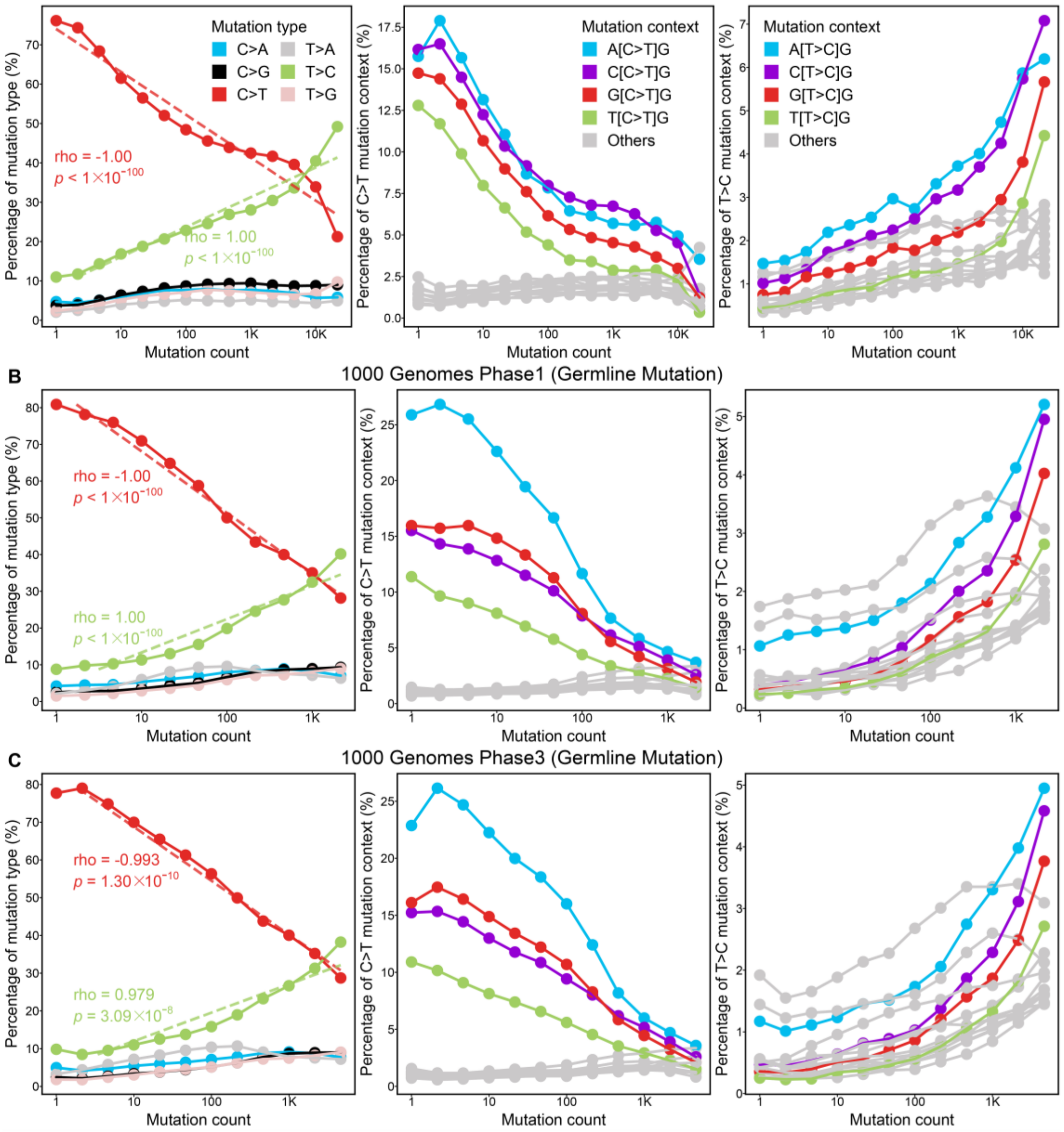


**Supplementary Figure S1. Frequency dependence of germline mutational spectra in various datasets.** The analysis was performed on germline mutations from (A) The Cancer Genome Atlas (TCGA), (B) 1000 Genomes Project Phase 1, and (C) 1000 Genomes Project Phase 3. Left panels show the relationship between the percentage of the six base substitution types (C>A, C>G, C>T, T>A, T>C, T>G) and the mutation count. The Spearman's rank correlation coefficient (rho) and p-value for the correlation between the percentage of C>T (red) and T>C (green) mutations and the mutation count are shown. Middle panels show the percentage of C>T mutations within various trinucleotide contexts, plotted against the mutation count. Right panels show the percentage of T>C mutations within various trinucleotide contexts, plotted against the mutation count. (B, C) use the same color legends as (A). The x-axis is represented on a logarithmic scale.

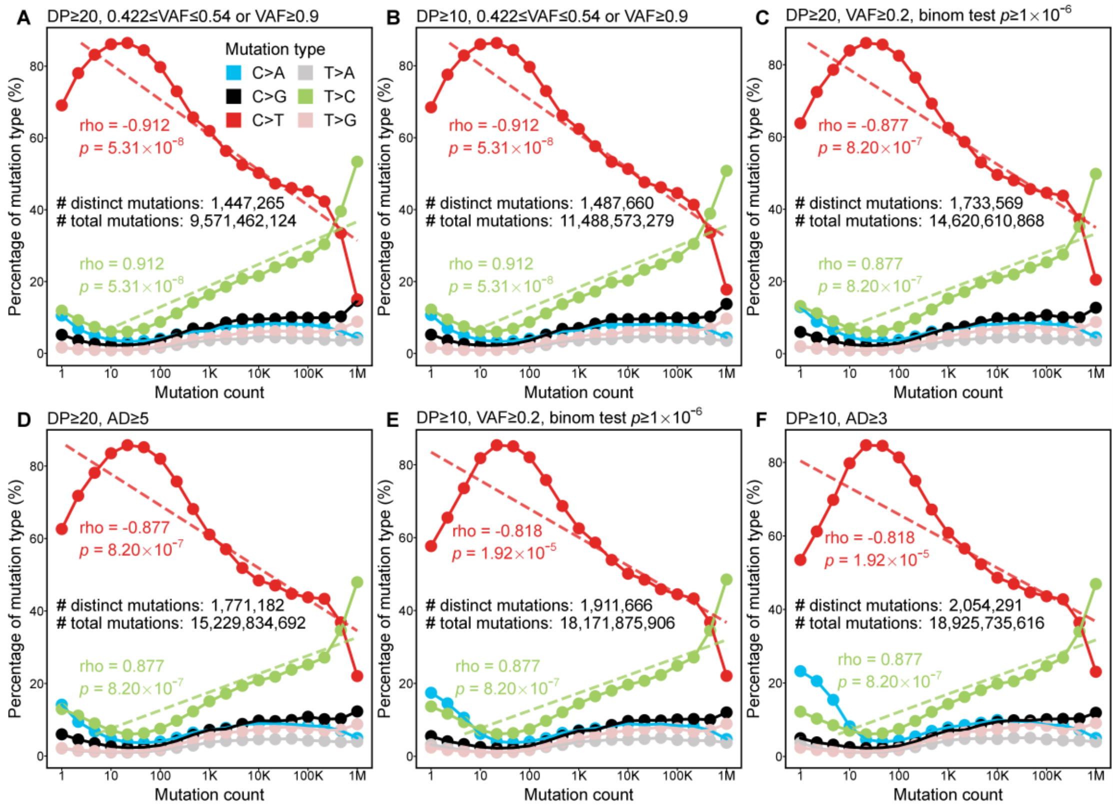


**Supplementary Figure S2. Robustness analysis of the relationship between germline mutational spectrum and mutation frequency.** The relationship between the percentage of the six base substitution types (C>A, C>G, C>T, T>A, T>C, T>G) and the mutation count, under six different mutation filtering criteria (A-F). The Spearman's rank correlation coefficient (rho) and p-value for the correlation between the percentage of C>T (red) and T>C (green) mutations and the mutation count are shown. (B-F) use the same color legend as (A). The x-axis is represented on a logarithmic scale. DP: total sequencing depth; VAF: variant allele frequency; AD: variant allele depth; binom test: binomial test between VAF and 50%.

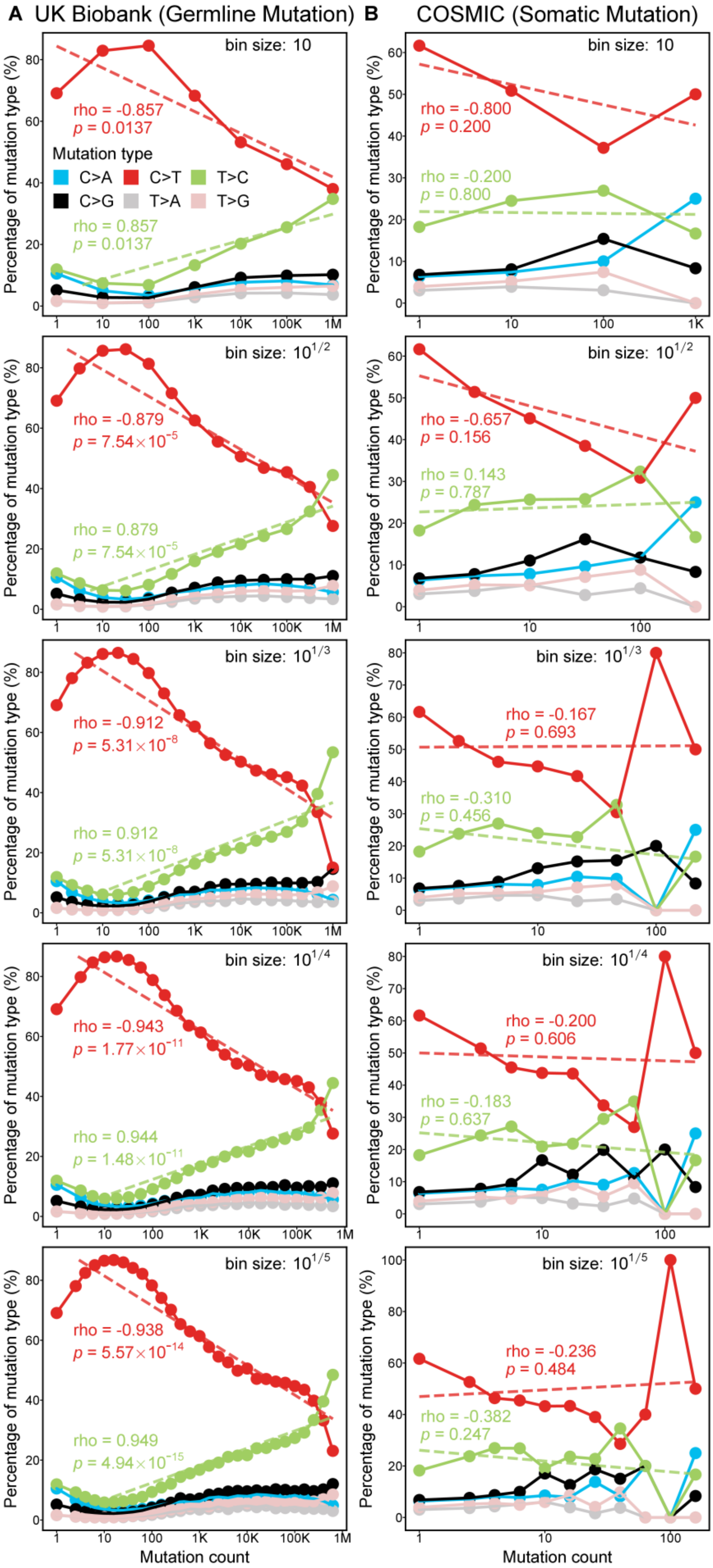

A UK Biobank (Germline Mutation)
B COSMIC (Somatic Mutation)
bin size: 10
rho = -0.857
p = 0.0137
Mutation type
C>A C>T T>C
C>G T>A T>G
rho = 0.857
p = 0.0137
rho = -0.800
p = 0.200
rho = -0.200
p = 0.800
bin size: 10^1/2
rho = -0.879
p = 7.54×10^-5
rho = 0.879
p = 7.54×10^-5
rho = -0.657
p = 0.156
rho = 0.143
p = 0.787
bin size: 10^1/3
rho = -0.912
p = 5.31×10^-8
rho = 0.912
p = 5.31×10^-8
rho = -0.167
p = 0.693
rho = -0.310
p = 0.456
bin size: 10^1/4
rho = -0.943
p = 1.77×10^-11
rho = 0.944
p = 1.48×10^-11
rho = -0.200
p = 0.606
rho = -0.183
p = 0.637
bin size: 10^1/5
rho = -0.938
p = 5.57×10^-14
rho = 0.949
p = 4.94×10^-15
rho = -0.236
p = 0.484
rho = -0.382
p = 0.247
Percentage of mutation type (%)
Mutation count

**Supplementary Figure S3. Frequency dependence of mutational spectra in germline vs. somatic mutations under different binning resolution.** (A) Analysis of germline mutations from the UKB. (B) Analysis of somatic mutations from the COSMIC database. Each panel show the relationship between the percentage of the six base substitution types (C>A, C>G, C>T, T>A, T>C, T>G) and the mutation count. Each row displays the results for a different logarithmic scale bin size of mutation count. The Spearman's rank correlation coefficient (rho) and p-value for the correlation between the percentage of C>T (red) and T>C (green) mutations and the mutation count are shown. All panels use the same color legend as the left-top one. The x-axis is represented on a logarithmic scale.
